# A novel capsid protein network allows the characteristic inner membrane structure of *Marseilleviridae* giant viruses

**DOI:** 10.1101/2021.02.03.428533

**Authors:** Akane Chihara, Raymond N. Burton-Smith, Naoko Kajimura, Kaoru Mitsuoka, Kenta Okamoto, Chihong Song, Kazuyoshi Murata

**Affiliations:** Department of Physiological Sciences, School of Life Science, The Graduate University for Advanced Studies (SOKENDAI), Okazaki, Aichi, Japan; National Institute for Physiological Sciences, National Institutes of Natural Sciences, Okazaki, Aichi, Japan; Exploratory Research Center on Life and Living Systems (ExCELLS), National Institutes of Natural Sciences, Okazaki, Aichi, Japan; Research Center for Ultra-High Voltage Electron Microscopy, Osaka University, Suita, Osaka, Japan; Program in Molecular Biophysics, Department of Cell and Molecular Biology, Uppsala University, Uppsala, Sweden

**Keywords:** Giant virus, NCLDV, capsid proteins, high-voltage electron microscope, cryo-electron microscopy, single particle analysis, homology model

## Abstract

*Marseilleviridae* is a family of the new order of giant viruses, which exhibit a characteristic inner membrane. Here, we investigated the entire structure of tokyovirus, a species of *Marseillevirus* at 7.7 Å resolution using 1 MV high-voltage cryo-EM and single particle analysis. The minor capsid lattice formed by five proteins, shows a novel structure compared to other icosahedral giant viruses. Under the minor capsid proteins, scaffold proteins connect two five-fold vertices and interact with the inner membrane. Previously reported giant viruses utilise “tape measure” proteins, proposed to control its capsid size, which could not be identified in tokyovirus, but scaffold proteins appear to perform a similar role. A density on top of the major capsid protein was identified, which suggested to be a 14kDa glycoprotein. Our observations suggest that the icosahedral particle of *Marseilleviridae* is constructed with a novel capsid protein network, which allows the characteristic inner membrane structure.

## Introduction

The “giant viruses” are exceptionally large viruses, larger than small bacteria^1^. They have a much larger genome than other viruses and contain many genes not found in other viruses^2^. All giant viruses belong to the superfamily of nucleo-cytoplasmic large DNA viruses (NCLDV). The NCLDVs are a phylogenetic group of viruses that possess double-stranded DNA and target varying host eukaryotes^3^. At present, the NCLDV group is said to be composed of eight families: *Poxviridae, Asfarviridae, Iridoviridae, Ascoviridae, Phycodnaviridae, Mimiviridae, Marseilleviridae*, and *Pithoviridae*^4^. In recent years, a variety of NCLDVs have been isolated and studied from around the world, and unclassified viruses such as pandoravirus and medusavirus also exist^5^. A new order, *Megavirales*, has been proposed based on the shared characteristics of these viruses^6^. More recently, *Mininucleoviridae* was proposed as a new family^7^.

*Marseilleviridae*, first isolated from amoebae in 2009, is a family of the new order of giant viruses, which exhibit a characteristic inner membrane structure extruded locally under the five-fold vertices of the icosahedral capsid^8^. Lausannevirus, senegalvirus, cannes 8 virus, tunisvirus, melbournevirus and marseillevirus belong to this family^9^. Although firstly isolated in amoebae, studies have also reported the presence of *Marseilleviridae* in humans^10, 11^, however there is controversy surrounding marseillevirus infections of humans as other studies have shown no evidence^12, 13^. *Marseilleviridae* has a particle size of ∼230 nm (250 nm maximum diameter, from opposing five-fold axes)^8^. Melbournevirus of the same family^8^ which has previously been analysed by cryo-electron microscopy (cryo-EM) single particle analysis (SPA) at 26 Å resolution, forms an icosahedral capsid with T = 309, and the lipid bilayer (inner membrane) inside the capsid covering the nucleoid. It was found that it has a characteristic structure that extrudes just below the five-fold axis, but it was not clear why such a structure was present. Tokyovirus is a giant virus of the family *Marseilleviridae* isolated from the eastern area of Tokyo^9^. In this study, we utilise tokyovirus to analyse the structural characteristics of the icosahedral *Marseilleviridae*.

Few studies have revealed the capsid structure of large icosahedral viruses in detail. An electron microscopy study half a century ago revealed that the NCLDV capsid is composed of 20 and 12 sets of trisymmetron and pentasymmetron, which are clusters of capsomer forming pseudo-hexamer, respectively^14^. In recent years, the structures of many proteins and viruses have been revealed with high resolution by using cryo-EM SPA for determination of overall structure^15^, dynamics^16, 17^ and assembly^18^. However, the NCLDVs present special challenges for high resolution cryo-EM simply from their size, which imposes hard limits on sample preparation, data acquisition and image reconstruction techniques. In 2019, *Paramecium bursaria* chlorella virus 1 (PBCV-1) of *Phycodnaviridae* and the structure of African swine fever virus (ASFV) of *Asfarviridae* at 3.5 Å resolution and 4.1 Å resolution, respectively, were reported by the use of cryo-EM SPA^19, 20^. The previous cryo-EM SPA of NCLDVs at high resolution have used microscopes with an accelerating voltage of 300 kV. However, these reports utilise a hybrid methodology, termed “block-based reconstruction”^21^ to achieve these resolutions which focusses on sub-sections (“blocks”) of the virus to permit localised defocus refinement, resulting in reconstructions of higher resolution. In these reports, it was shown that the capsids of PBCV-1 and ASFV are composed of one “major” capsid protein (MCP) and a combined fourteen kinds (PBCV-1) or four kinds (ASFV) of “minor” capsid protein (mCP). It has been reported in other NCLDVs^19, 20^ that the MCP has two “jelly roll” motifs. The jelly roll motif is composed of eight β strands^22^. The mCP forms a hexagonal network to fill the space between the MCPs, which likely contributes toward keeping the capsid structure stable. Further, it was reported that PBCV-1^20^ and ASFV^19^ possess an mCP called “tape-measure” protein. The tape-measure protein is a long filamentous protein that extends from a pentasymmetron to an adjacent pentasymmetron along the trisymmetron edge. It was proposed that the length of this tape-measure protein determines the capsid size^23^.

Herein we focus on the single particle reconstruction of the whole tokyovirus using 1 MV cryo-EM. The 1 MV high-voltage electron microscope (HVEM) was originally developed to extend attainable resolution using shorter electron wavelengths^24^. However, major usage is currently focussed on thick specimens, which lower acceleration voltages are unable to penetrate^25^. High voltage cryo-electron microscopy (cryo-HVEM) on biological samples has not yet been reported for single particle analysis, and only a few examples using tomography have been reported^26, 27^, ^28^. In the case of a thick sample (for example, a 250 nm diameter virus) an internal focus shift occurs due to the influence of the depth of field, imposing a hard limit on attainable resolution^29^. However, by increasing the accelerating voltage (and therefore shortening the wavelength of the emitted electrons) the depth of field is increased, and it is possible to improve the (electron)-optical conditions in a thick sample. The resolution achievable at a given acceleration voltage can be calculated by the following formula

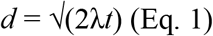

Where *d* is the resolution at which all points within a given thickness can be considered equally focussed, *t* is the sample thickness (in this case, particle diameter), and λ is the electron wavelength^30^. It should be noted that for high symmetry specimens, such as icosahedral viruses, sample thickness (or depth) is functionally equivalent to particle width. Previous discussions^31, 32^, ^33^ cover the equations for calculating electron wavelengths and accounting for relativistic effects at given acceleration voltages if one wishes to further explore the mathematics. Using Eq. 1, for a 250 nm thick sample, at an acceleration voltage of 300 kV, the electron wavelength is 1.97 pm, so the theoretical resolution limit is ∼9.9 Å (Fig. 1). Extending this, at an accelerating voltage of 1 MV, the electron wavelength is 0.87 pm, so the theoretical resolution limit is calculated to improve to 6.6 Å. From the above, we attempted to clarify the structure of the entire giant virus particle with higher resolution using a 1 MV cryo-HVEM rather than using the block-based approach.

**Fig. 1.**
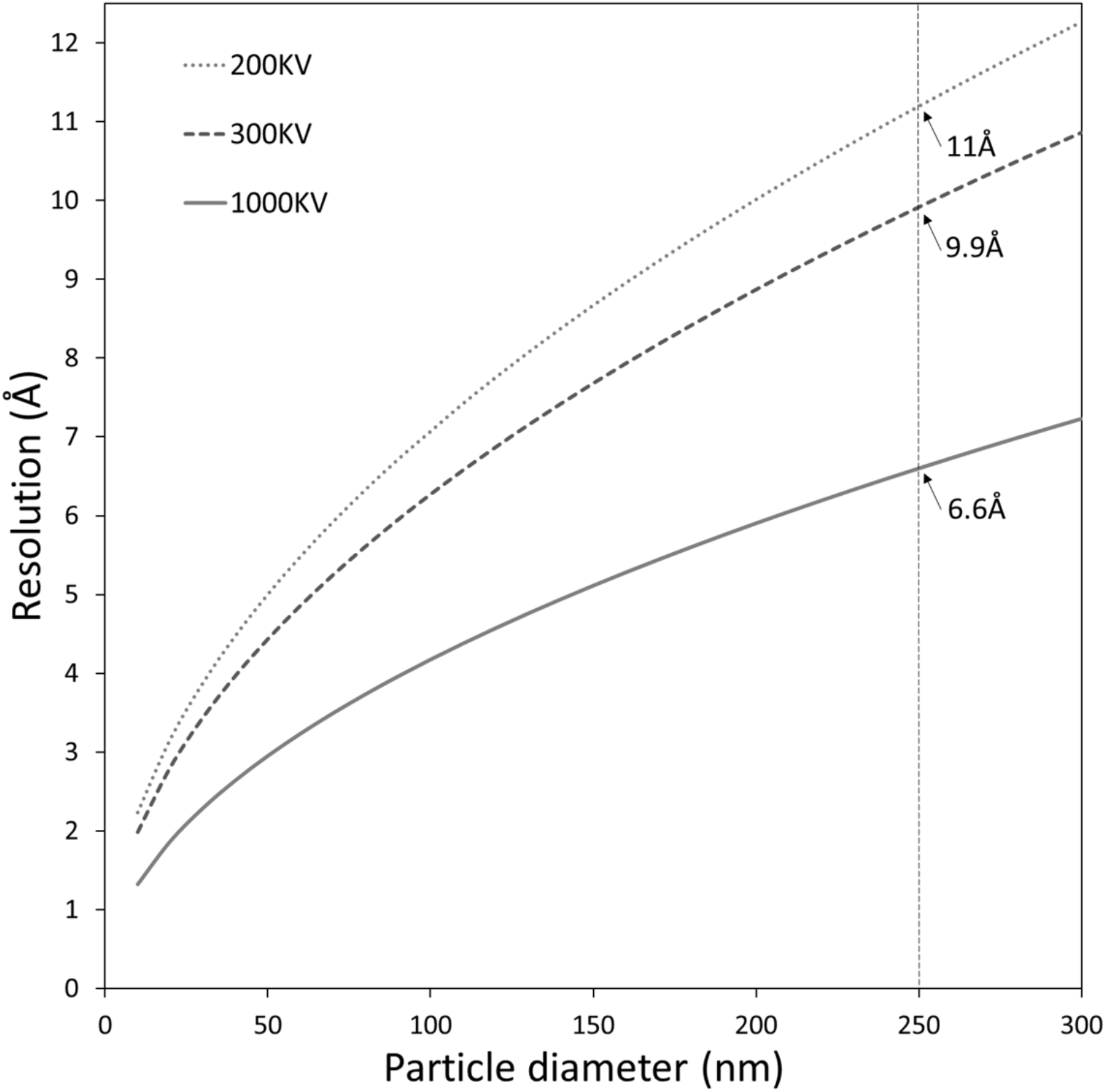
The depth of field effect in cryo-EM. Theoretical resolution limits caused by the size of particles at 200, 300 and 1,000 kV, from 100-3000 Å. Dashed vertical line drawn at approximate maximum diameter of tokyovirus. Theoretical resolution limit at the approximate maximum diameter of tokyovirus (250 nm) is shown to one decimal place.

In this study, we first estimate the theoretical resolution limit in thick specimens imposed by using a 1 MV cryo-HVEM and use a platinum-iridium (PtIr) film to examine the performance of the microscope itself. Following this, three-dimensional (3D) reconstruction of tokyovirus was performed using data acquired with the cryo-HVEM. We achieve close to the theoretical resolution limit. From the obtained 3D reconstruction, we show that the capsid of tokyovirus is composed of MCP, eight distinct types of mCP, scaffold protein (ScP), and penton protein, with an oddly extruded inner membrane containing the viral genome. ScP exists between the mCP layer and the internal membrane, and a part of the ScPs is bound to the internal capsid proteins immediately below the five-fold axis, which is characteristic of *Marseilleviridae*. Since this is a structure not reported in other NCLDV, we hypothesise that it is important in the maintenance of the capsid of marseilleviruses. Our results pave the way for a greater understanding of this family of NCLDV, in addition to the SPA of gigantic biological specimens.

## Results

### Performance of the 1 MV HVEM

While the theoretical resolution limit imposed by depth-of-field is improved by moving to higher acceleration voltages (Fig. 1)^29, 30^, it is of little practical use if other factors are providing hard limits to attainable resolution. To this end, we examined the performance of the 1 MV cryo-HVEM (JEOL JEM-1000EES) using a Pt-Ir film and performed rotational averaging on a calculated power spectrum (Fig. 2a). In the 1D intensity plot, Thon rings are clearly visible to 1.81 Å (Fig. 2b). Comparing the performance of the 1 MV cryo-HVEM equipped with a LaB^6^ electron gun and Gatan K2 Summit direct detector against a high performance 300 kV microscope using a thermal field emission electron gun (in this case a FEI Titan Krios was used) is also of great interest (Supplementary Fig. 1). The Modulation Transfer Function (MTF) of the 1 MV microscope is superior in counting mode across the entire spatial frequency range, although in super resolution mode it suffers slightly below 0.25 Nyquist frequency (Supplementary Fig. 1a). In addition, the MTF of the cryo-HVEM shows a 5% reduction in performance with super resolution acquisition at the lower special frequencies compared to the 300 kV cryo-EM. Detective Quantum Efficiency (DQE) is similarly superior for the 1 MV microscope in counting mode, but to our surprise performance in super resolution mode shows a significant reduction in performance in both 1 MV and 300 kV microscopes. Furthermore, DQE of the cryo-HVEM show as much as a 40% drop in performance in the lower special frequencies compared to the 300 kV cryo-EM. Beyond 0.6 Nyquist frequency, however, the two are comparable (Supplementary Fig. 1b). In this study, super resolution mode was used to increase sampling efficiency for the large-size virus particles (Table 1).

**Fig. 2.**
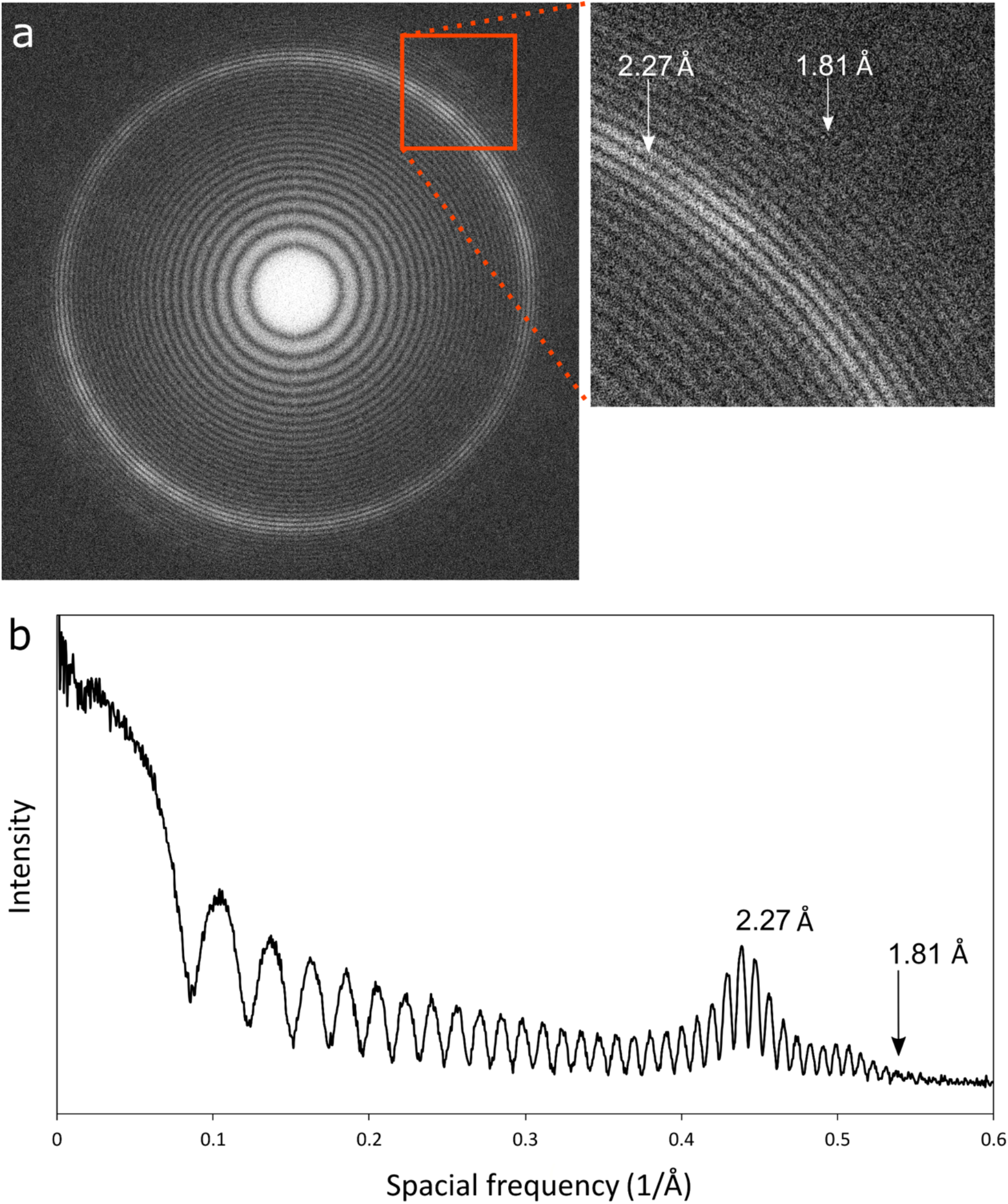
Performance of the cryo-HVEM (JEOL JEM-1000EES) equipped with a Gatan K2 Summit direct electron detector using a Pt-Ir standard sample. a) Power spectrum showing clear Thon rings. b) A focussed view of the Thon rings. c) 1D plot of the power spectrum calculated with the “Radial Profile Extended” ^66^ plugin and Fiji ^65^.

**Table 1:**
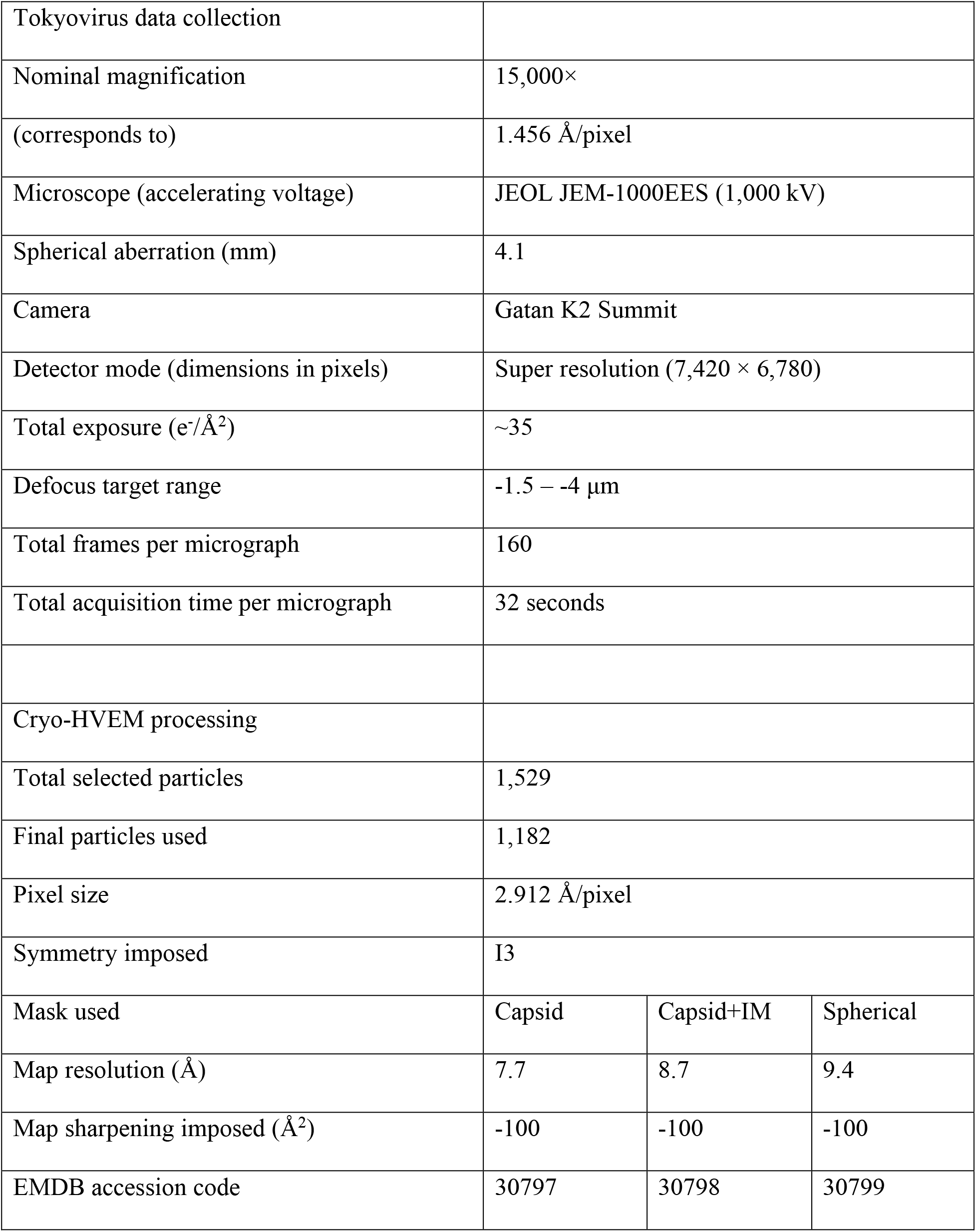
Data collection information

### Overall structure of Tokyovirus

Fig. 3 shows the overall structure of tokyovirus reconstructed using RELION 3.1 with icosahedral symmetry imposed. Tokyovirus has an outer capsid shell comprised of MCPs arranged in an icosahedral form with T=309 (h=7, k=13) (Fig. 3a). Immediately below the outer shell are layers of mCPs and ScPs in that order, and the innermost membrane stores viral nucleic acids (Fig. 3b-d), while the relatively large mCPs (arrow in Fig. 3c) directly interact with the inner membrane extrusion under the 5-fold vertices. In the extrusion of inner membrane, membrane forms multilayers (asterisk in Fig. 3c). Compare to MCP and mCP, density of ScP is smeared, showing a structural flexibility of the component (Fig. 3c, d). Fig. 3e shows the estimated local resolution of the capsid layer of Tokyovirus, sliced for internal visualisation. The highest resolution was shown in the interface between MCP and mCP. Each part has a characteristic structure, which can be better viewed by careful filtering and segmentation of the reconstruction (Fig. 4). MCP covers whole surface of the viral capsid except for the 5-fold vertices (lavender in Fig. 4), where the penton protein is plugged the hole in the vertices (horn in Fig. 4b mCP). mCP is roughly distinguished by three components, where components under pentasymmetron, trisymmetron, and border of trisymmetron (blue in Fig. 4). The ScP array is located further inside the virus between the mCP layer and internal membrane (yellow in Fig. 4). Unlike the other mCPs, the ScPs form an anti-parallel chained array between pentasymmetrons along each two-fold axis, thus bounding each trisymmetron (yellow in Figs. 4, 5). The internal membrane has a characteristic structure that extrudes outwards below each five-fold axis (Fig. 3b-c, grey in Fig. 4), which appears to be formed by a frame of ScPs.

**Fig. 3.**
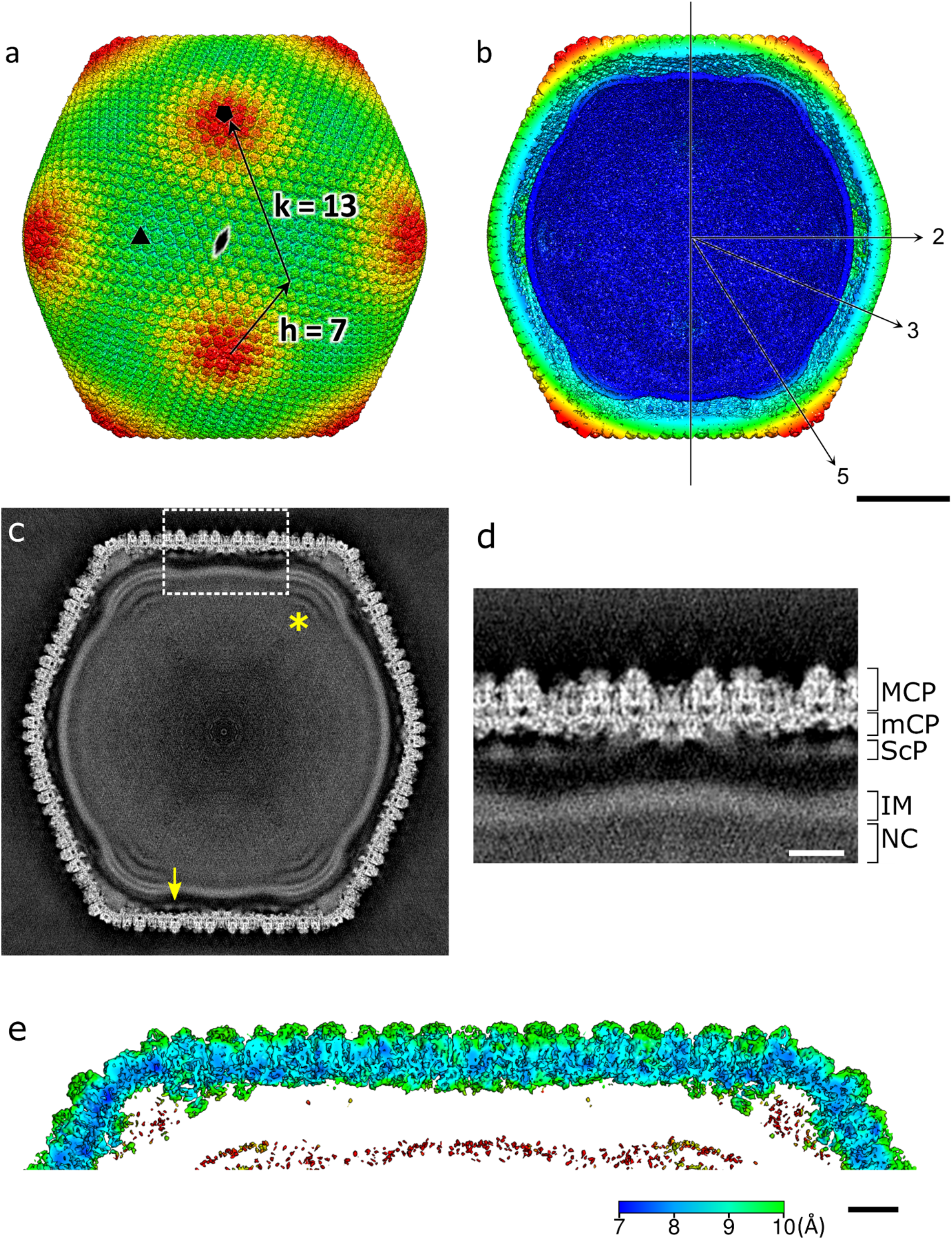
SPA 3D reconstruction of Tokyovirus at 7.7 Å. a) Tokyovirus, coloured by radius, and showing the five-fold (black pentagon), three-fold (black triangle) and two-fold (black double-teardrop) symmetry axes. b) Tokyovirus cross-section, with symmetry axes shown with arrows. (a, b) are coloured by radius in UCSF Chimera with the following parameters: blue, 910 Å; turquoise, 1010 Å; green, 1080 Å; yellow, 1125 Å; red, 1200 Å. c) A central slice of Tokyovirus 3D reconstruction. d) Focussed view of the marked box in (c). showing delineation between major capsid protein (MCP), minor capsid protein (mCP), scaffold proteins (ScP), inner membrane (IM) and nucleocapsid (NC). e) local resolution of the capsid, estimated by the *blocres* module of Bsoft ^49^, focussing on the upper capsid edge, sliced to permit visualisation of internal density. Scale bars: a-c 50 nm, d-e 10 nm.

**Fig. 4.**
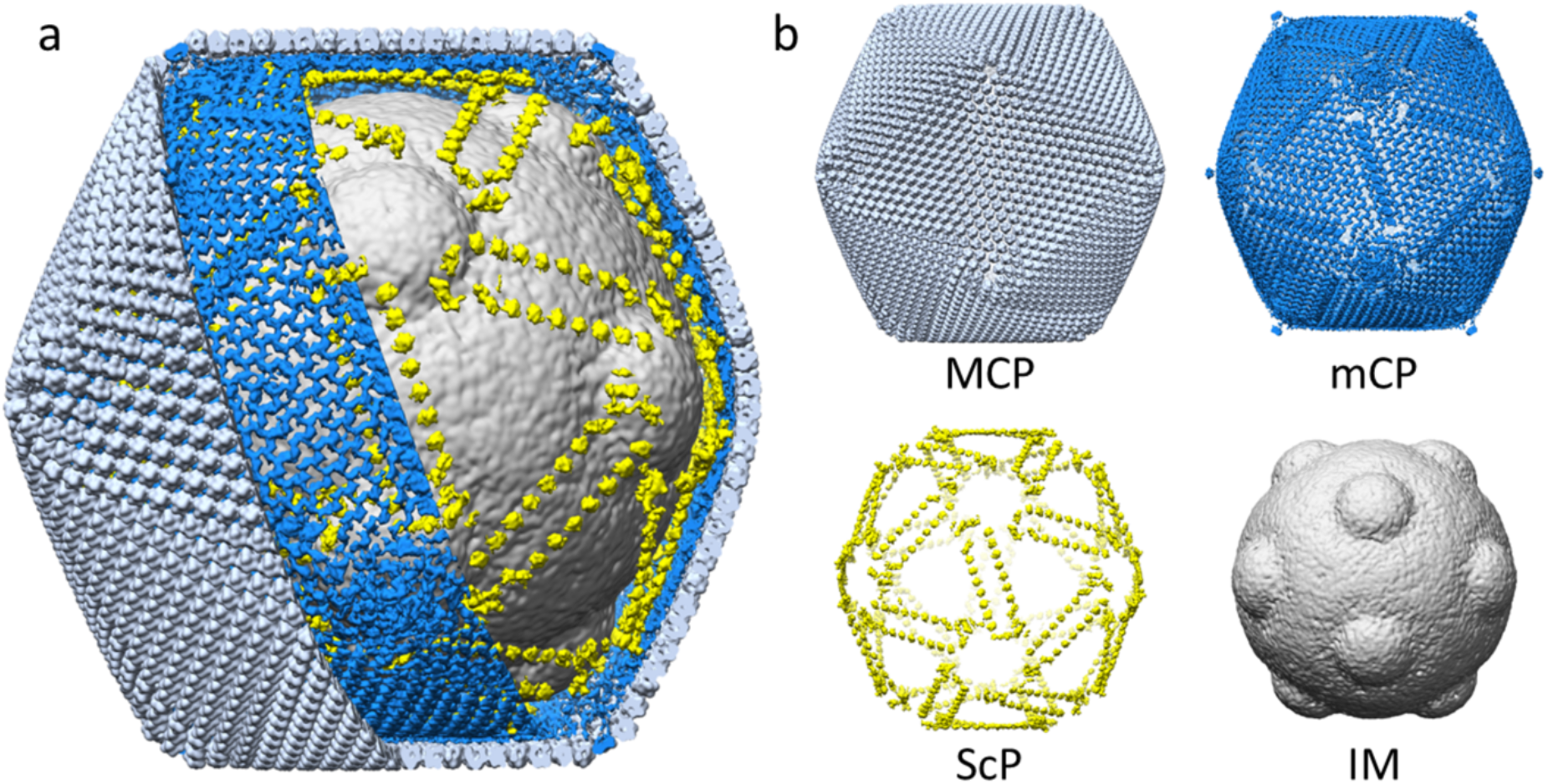
Segmentation of tokyovirus structure. a) The complete tokyovirus virion cut out to show each component. b) Individual components of the virion; major capsid protein (MCP), minor capsid protein (mCP), scaffold protein array (ScP) and inner membrane (IM). Individual segments (e.g.: inner membrane) are low-pass-filtered to improve visualisation clarity.

**Fig. 5.**
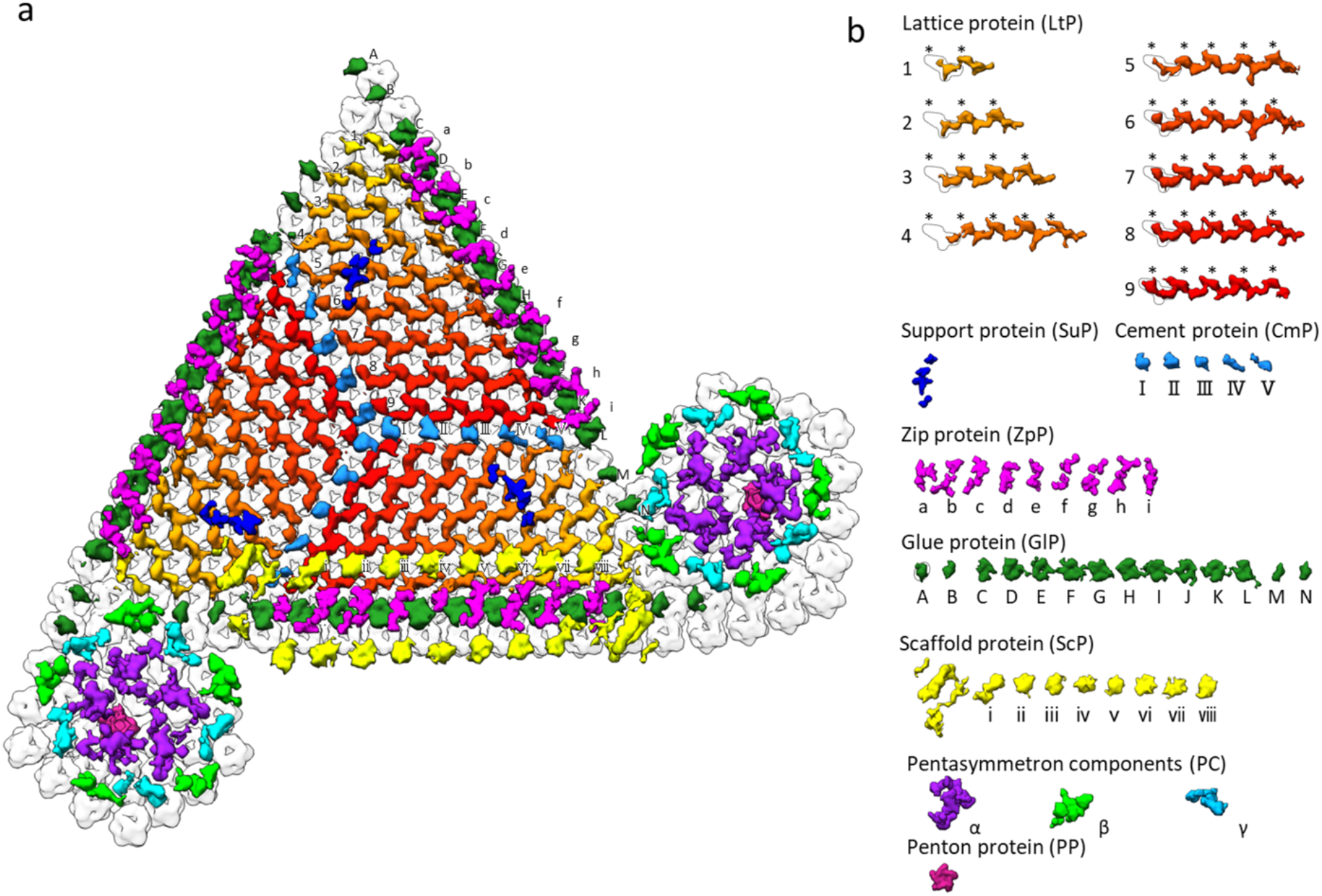
Focussed segmentation of minor capsid proteins, scaffold protein (ScP) and penton protein (PP) of tokyovirus. mCPs were classified into 8 components consisted of 9 lattice protein repeats (LtPs 1-9), 1 support protein (SuP), 5 cement proteins (CmP I-V), 9 zip proteins (ZpP a-i), 14 glue proteins (GlP A-N), and 3 pentasymmetron components (PC-α, β, and γ).

### Minor capsid proteins (mCPs)

We classified the mCPs into eight types of proteins based on their structural features and arrangements (Fig. 5). They will be referred to as Lattice protein (LtP) (orange to red in Fig. 5), Support protein (SuP) (royal blue in Fig. 5), Cement protein (CmP) (sky blue in Fig. 5), Zip protein (ZpP) (pink in Fig. 5), and Glue protein (GlP) (emerald green in Fig. 5), three pentasymmetron components (PC-α, β, and γ) (purple, light green, and cyan in Fig. 5), respectively. The mesh-shaped triangle formed by mCPs is composed of three trapezoidal units consisting of these five kinds of proteins (LtP, SuP, CmP, ZpP, GlP) each rotated by 120° about the three-fold rotation axes. These proteins are connected to PC-β and PC-γ at the edge of the triangle.

Lattice protein (LtP) forms a wavy structure along the gap of MCP by a repeating “running dog” shape structure (orange to red in Fig. 5). This waveform structure consists of nine columns in one unit (1-9 in Fig. 5), and the number of repetitions of the structure changes according to the number of MCPs present above each. It forms the base of the trapezoidal unit. The electron density of the trisymmetron interface point at the end of the waveform (dotted circles in LtP in Fig. 5) cannot be clearly confirmed where it binds to the adjacent ZpP. Under the network structure formed by LtP, the SuP creates a bridge from LtP4 to LtP6, further appearing to perform a role in strengthening the mCP array by forming a bridge between the ScP/ZpP/GlP and the parallel CmP array (Fig. 5, Fig. 6a, b). Within the same trisymmetron, the CmP connect the three trapezoidal units by interfacing the ends of LtP5 to LtP9, extending from the three-fold axis toward the associated pentasymmetron, terminating at the ScP edge.

**Fig. 6.**
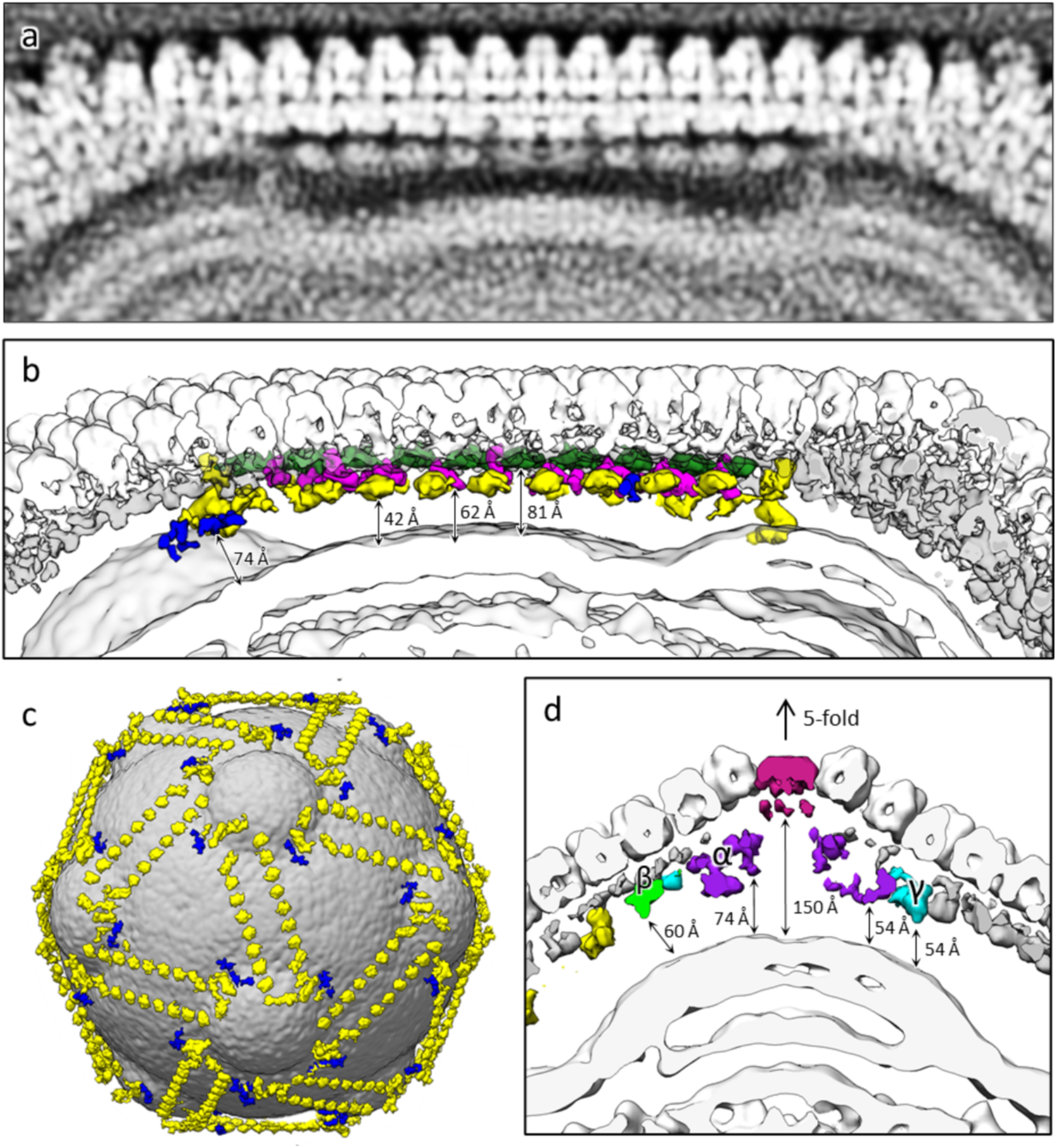
Examination of the scaffold protein network between the inner membrane and capsid shell. a) Slab density of the tokyovirus showing the innermost layer of the mCP network. b) Isosurface view of (a) coloured as in Fig. 5 (GlP; green, ZpP; hot pink, ScP; yellow, SuP; royal blue) and with distances from the inner membrane to the components indicated by arrows. c) The tokyovirus inner membrane with ScP and SuP network overlaid. d) Focussing on the pentasymmetron at the five-fold axis, representing distances from the inner membrane extrusion to PC-α, β, γ and the PP.

Two proteins, ZpP and GlP, are involved in connecting adjacent trisymmetrons and run directly above the ScP array. GlP (emerald green in Fig. 5) is present at the boundary between adjacent trisymmetrons and appears to glue them. ZpP “fills the gaps” between LtPs at the trisymmetron interfaces, interlocking with GlP like the teeth of a zipper. There are 14 GlPs (A to N) and 9 ZpPs between trisymmetrons (Fig. 5). C-L of GlPs have a flat circular shape, but the termini GlPs (A, B, M, N) appear roughly semicircular.

Scaffold protein (ScP) is present further inside the mCP layer (yellow in Fig. 5, Fig. 6a, b), forming a framework surrounding the inner membrane. It interacts with both the inner membrane and the capsid (MCPs and mCPs) (Fig. 6a, b). ScP has a large U-shaped structure at the end, from which 8 round structures are arranged along the edge of trisymmetron (Figs. 5 and 6c). Two ScPs are anti-parallelly connected with the head and tail, and both edges are located at the apex of the pentagon of pentasymmetron (Fig. 6c). The adjacent ScP terminal pairs are further connected with SuP (royal blue in Fig. 6c).

### Penton protein and pentasymmetron

The penton density (central region, Supplementary Fig. 2a) is similar to that of PBCV-1 (Supplementary Fig. 2b), in that it possesses a “cap” region which aligns with the MCP layer and a lower region which aligns with the mCP layer. ASFV, by comparison, is missing this lower region (Supplementary Fig. 2c). We fitted the PDB model of the PBCV-1 penton (Supplementary Fig. 2e) to the upper region of the tokyovirus penton (Supplementary Fig. 2d).

In ASFV, the insertion domain is extrinsic of the MCP interface (Supplementary Fig. 2f), while neither tokyovirus nor PBCV-1 exhibit clear density for this domain. The penton density including the lower region itself has a distance to the inner membrane extrusion of ∼150 Å (Fig. 6d).

The three PC (α, β, and γ) proteins maintain a similar distance from the IM extrusion to PCs (54-60 Å) (Fig. 6d), which is also similar to that of the ScP array along the two-fold axes (42 Å) (Fig. 6b). PC-α is the largest, interacting with each other and the PP itself (Fig. 5a). Pairs of PC-β and PC-γ surround this group of PC-α (Fig. 5a). PC-β appears to interact primarily with a single PC-α and acting as terminal for the GlP. PC-γ appears to interact with two neighbouring copies of PC-α. These proteins function to support MCP array in the pentasymmetron, maintain mCP network with those under trisymmetron, and interact with the extrusion of the inner membrane under the five-fold vertices.

### Inner membrane (ScP interaction)

The internal structure of *Marseilleviridae* possesses a very characteristic deformation of the internal membrane at the five-fold axes^8^. Tokyovirus also possesses this inner membrane structure. Within this deformation, there are multiple layers (Fig. 3c, Fig. 6a-b, d). The extruded membrane region and ScP appear to interact. It is unclear whether the inner membrane deforms to fill the cavity left by the ScP array, or whether the ScP array plays a role in distorting the membrane into this shape at the five-fold axes (Fig. 6c, d). This needs further study in the future.

### Major capsid protein

A BLAST search^34^ was performed using the proposed MCP sequence from the draft genome of tokyovirus^9^ (Table 2). The MCP of Tokyovirus scores highest *versus* iridovirus (PDBID: 6OJN)^35^ and PBCV-1 (PDBID: 5TIP)^36^, with high coverage and moderate homology, although both comparisons of conserved sequences fall below the “identities” that allows direct and safe comparisons^37^. The three respective MCP amino acid sequences were aligned by PROMALS3D^38^. The secondary structure of tokyovirus was predicted by PSIPRED^39^ and the secondary structure of iridovirus and PBCV-1 in the PDB model. As a result, tokyovirus MCP was predicted to possess two sets of eight β strands (B1 to I2) that form a jelly roll motif like other NCLDVs (Supplementary Fig. 3)^6^.

**Table 2:** Results of BLAST comparison of Tokyovirus MCP versus other NCLDVs

| Virus | PDB ID | a.a. | BLAST search |  |  |  |
| --- | --- | --- | --- | --- | --- | --- |
|  |  |  | E-value | Score | Query Cover (%) | Identities (%) |
| Iridovirus | 6OJN | 463 | 1.00E-105 | 310 | 99 | 38.21 |
| PBCV-1 | 5TIP | 436 | 5.00E-17 | 68.9 | 96 | 19.92 |
| Faustovirus | 5J7O | 645 | 0.016 | 23.9 | 30 | 37.5 |
| Phage PM2 | 2VVF | 269 | 0.043 | 21.2 | 14 | 26.67 |
| ASFV | 6KU9 | 693 | 0.069 | 21.9 | 38 | 12 |
| Sputnik virus | 3J26 | 508 | 0.48 | 18.9 | 27 | 36 |
| PRD-1 | 1CJD | 394 | 0.93 | 17.3 | 16 | 37.04 |
| Vaccinia virus | 2YGB | 569 | 2.4 | 16.5 | 14 | 29.63 |
| Phage FLiP | 5OAC | 310 | 2.7 | 15.4 | 15 | 42.86 |
| Adenovirus | 6B1T | 952 | 3 | 16.9 | 16 | 21.43 |
| STIV | 2BBD | 350 | 4.4 | 15 | 13 | 22.73 |
| Mavirus | 6G45 | 610 | 4.8 | 15.8 | 12 | 25.93 |

In the MCP amino acid sequence of each virus, there is a large difference in loop length between β strands forming the jelly roll motif (Supplementary Fig. 4). To investigate how the difference in loop length appears in the difference in the structure of MCP, a homology model of tokyovirus MCP was generated based on PBCV-1, using the SWISSMODEL server^40^, and compared to both PBCV-1 and iridovirus (Supplementary Fig. 4). This was used despite the relatively low homology as it is the only MCP structure of an NCLDV built from a density map rather than *in silico* (as iridovirus). This homology model was further optimised to fit the MCP density using COOT^41^ and PHENIX^42^. In tokyovirus MCP, the loops connecting the upper part of the jelly roll 1 (JR1) and JR2 are longer than that of PBCV-1. The HI1 loop is particularly extended in tokyovirus (Supplementary Figs. 3 and 4) the 23-residue sequence is longer than that of PBCV-1. The MCP unit from a three-fold axis of tokyovirus was extracted (Fig. 7a), and the homology model built into the 7.7 Å resolution map (Fig. 7b, Supplementary Fig. 4a). The JR1 loops, which constitutes the upper part of the MCP including the HI1 loop, coincides with the upper part density of the tokyovirus MCP (Fig. 7b), but does not fill the entire cap density of the tokyovirus MCP (Fig. 7b, c). The density in the tokyovirus MCP which is not filled by the fitted homology model has been highlighted in orange (Fig. 7c). The HI1 loop is significantly extended beyond that of PBCV-1 and marginally longer than that of iridovirus (Supplementary Fig. 4).

**Fig. 7.**
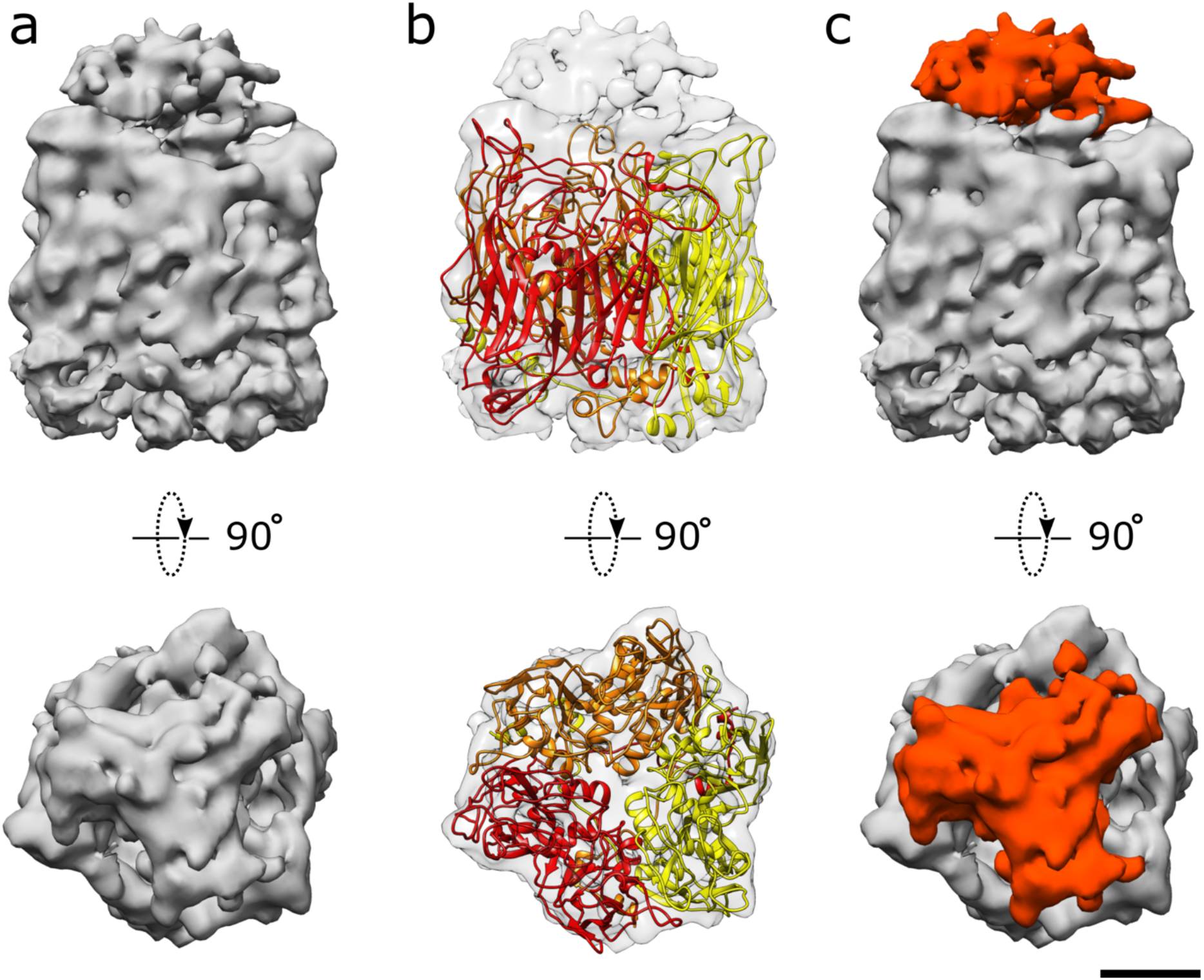
Cryo-EM map and fitted homology model of tokyovirus MCP. The homology model was generated by SWISS-MODEL and adjusted to the volume with COOT and PHENIX. a) Side and top views of the extracted cryo-EM map of MCP. b) Side and top views of the extracted MCP cryo-EM map with homology model fitted. c) Side and top views of the extracted MCP cryo-EM map with the empty cap region coloured in orange. Scale bar equals 2 nm.

The DE1 and FG1 loops are longer than those of PBCV-1 and of similar length to those of iridovirus. Conversely, the DE2 loop in tokyovirus is truncated compared to those of both PBCV-1 and iridovirus. Other loops are of comparable length.

The MCP density has a visible cap (Fig. 7c) which does not correspond to any part of the fitted homology model or fitted model of PBCV-1 MCP. The symmetric nature of this cap implies the presence of protein rather than disordered post-translational glycosylation. This is supported by Periodic acid Schiff (PAS) staining^43^ of SDS-PAGE separated tokyovirus proteins, which do not show any glycosylated proteins at the molecular weight of the MCP. However, a band is evident at ∼14kDa (Supplementary Fig. 5), indicating that there is likely to be another protein which has been glycosylated interacting with the top of the MCP.

## Discussion

Here we show the structure of Tokyovirus as a complete viral particle, using 1 MV accelerating voltage cryo-HVEM to overcome some of the resolution limit imposed by the depth of field effect on exceptionally large particles. Thus far, this is the highest resolution reconstruction of a giant virus larger than 200 nm without utilising the block-based reconstruction^21^ technique or similar methods^19^. At present with cryo-HVEM, micrographs must be manually collected. If automated acquisition were possible on the HVEM in the future, it is likely the resolution would further improve^44^. Fig. 1 demonstrates the theoretical advantage in resolution gained using greater accelerating voltages. Using 300 kV cryo-EM shows a maximum theoretical resolution of 9.9 Å. However, using 1 MV cryo-HVEM improves this to a resolution (6.6 Å) where some secondary structure should be discernible^45^ without the use of block-based reconstruction methods. In addition, the performance of the system was confirmed with Thon rings of Pt-Ir standard extending beyond 2 Å (Fig. 2). This demonstrates that the microscope would not be a limiting factor in resolutions achieved with the giant virus particles. Use of the cryo-HVEM imposed its own limits, however, requiring fully manual acquisition and acquisition times exceeding 30 seconds per micrograph. Further, some elements of general-purpose cryo-EM SPA image processing do not support HVEM accelerating voltages; Gctf^46^ was not used as it does not support 1 MV data, and likewise, dose weighting of 1 MV data is unsupported in both MotionCor2^47^ and the RELION implementation^48^. Fortunately, CTFFIND does support 1 MV data, which was used. We were unable to use dose weighting, although this is somewhat offset by higher acceleration voltages causing less radical damage. Particle polishing also provides little benefit, which may be caused by the large size of the viruses on the micrograph where distortions across the micrograph potentially cause virus deformation variation per frame. The size of the virus imposed other limits; requiring careful balance between lower magnification (to increase particles collected with manual collection and to reduce box size) and a high enough magnification to achieve worthwhile resolutions in reconstructions. Tokyovirus is so large that even at relatively low magnifications, there were at best around 12 virus particles per micrograph (often ∼6 virus particles, Supplementary Fig. 6a) which can be utilised for 2D classification, i.e.: do not include the edge of the micrograph. It was for this reason that “super resolution” mode in K2 Summit camera was used for data acquisition, though the detector performance is limited (Supplementary Fig. 1). The recent report regarding ASFV^19^ attained fewer particles per micrograph but were able to offset this using automated acquisition. As with other NCLDVs, tokyovirus presents a complex structure of multiple layers surrounding the central DNA core. For this reason, we utilised different masks for calculating the “gold standard” FSC during the post processing step to focus on different regions of the virus. Local resolution estimation is unaffected by these masks which affect the global FSC estimated resolution, (Fig. 3e, Supplementary Fig. 6h) and was calculated using *blocres* from Bsoft^49, 50^ utilising independent half-maps without masking. These software factors do provide some limitations for 1 MV cryo-HVEM for biological samples at present.

Ultimately, using cryo-HVEM provides benefits for larger particles. The decreased focus gradient across a particle aids high resolution whole particle reconstruction. Sample damage from the electron beam is also decreased, which is of particular importance with biological samples. Furthermore, increased beam penetration permits improved visualisation of internal structure, as used in cryo-tomography with pithovirus, an amphora-shaped giant virus^28^. Beyond that of more widely available 300 kV cryo-electron microscopes, it permits study of finer details of even larger viruses, which lower acceleration voltages are unable to penetrate. Finally, the reconstruction methodology is simpler than that of block-based reconstruction. However, we are applying block-based reconstruction methods currently to attempt to increase resolution further. As one purpose of block-based reconstruction is to overcome the defocus gradient which imposes such a hard limit at 300 kV, block-based reconstruction may yield little improvement at higher acceleration voltages. A second reason for using block-based reconstruction is to overcome minor distortions of very-large symmetric particles, yielding improved resolutions. However, if a particle is truly symmetric, then it may be better to consider distorted particles as damaged, rather than desirable to use.

The T=309 (*h*=7, *k*=13) capsid is clearly displayed (Fig. 3). With this structure we can newly identify an intermediate “scaffold” array between the viral capsid and the inner membrane. There do not appear to be any directly analogous densities to the “tape measure” protein of PBCV-1 and AFSV, the long, filamentous mCP named P2 in PBCV-1 and M1249L in AFSV, indicating that tokyovirus capsid construction occurs through a different mechanism. This lack of the “tape measure” to control capsid extension during virus construction, and presence of this scaffold network may indicate that the scaffold is responsible for imposing restraints on the capsid dimensions. Greater clarity of these flexible scaffold proteins will be required to further elucidate their interactions in more detail and perhaps shed more light on their potential role in construction.

The trapezoidal array of LtPs in tokyovirus do not form a single underlayer beneath the MCP in the manner of mCPs of PBCV-1 and ASFV, instead relying on another cement protein (CmP) which extends outwards from the center of the trisymmetron to connect the three rotated trapezoids (Fig. 5). Two proteins (GlP and ZpP) are present at the the trisymmetron edges rather than a single protein (P11) as in PBCV-1. An as-yet-unidentified protein of SuP also interacts with each trapezoidal lattice which does not appear to be present in reconstructions from PBCV-1 and ASFV.

Thus far, all NCLDVs appear to share a conserved MCP secondary structure featuring twinned “jelly roll” structures^19^ and the predicted structure of tokyovirus MCP follows this (Supplementary Fig. 3). However, the lengths of the unstructured loops between the β-sheets which form the two jelly roll folds varies (Supplementary Fig. 4). The HI1 loop of tokyovirus is extended, potentially playing a role in coordinating the “cap” protein. BLAST comparisons between the NCLDV MCPs show poor coverage versus the tokyovirus MCP in all cases except iridovirus and PBCV-1 (Table 2), while “identities” remains low for all NCLDV BLAST comparisons (Table 2, Supplementary Fig. 3). As a result, rigid body fitting of PBCV-1 MCP into a tokyovirus MCP density showed a poor fit.

To build the MCP homology model, we extracted the central MCP from a trisymmetron and the homology model rigid-body-fitted into the density was refined using COOT and PHENIX software (Fig. 7). An empty “cap” region was present on each MCP density when the MCP homology model was fitted to it, although thus far we have been unable to clarify this region sufficiently for model fitting. As such, we were initially unsure whether this density is caused by another, smaller protein interacting with the MCP or post-translational modification of the MCP. We identified a candidate of the small “cap” protein, as Periodic acid Schiff (PAS) staining was used on SDS-PAGE of purified tokyovirus particles to identify potential glycoproteins. This showed a protein running at ∼14kDa (Supplementary Fig. 5). In the case of PBCV-1, sugar chains are directly bound on the MCP^36^, while in the case of ASFV, there is a second membrane external of the capsid itself^19^ so capsid glycosylation is less important. This may suggest a different function of the glycosyl sugar chain in *Marseilleviridae*. We previously found some species of marseillevirus show that virus-infected host amoebae demonstrate a bunch formation^51^. Simultaneously, newly borne viruses were stuck and aligned on the host cell surface in the process. Another member of the *Mimiviridae*, tupanvirus, has evidence suggesting this similar mechanism^52^. The glycoprotein may play a role in this bunch formation, potentially to increase speed of transmission, but it needs further study to clarify the function.

The tokyovirus and PBCV-1 pentons are similar in depth (Supplementary Fig. 2a, b). The tokyovirus penton protein density is fitted with that of the cryo-EM-derived PDB model of PBCV-1 (PDBID: 6NCL) (Supplementary Fig. 2d) with density present for the penton protein and an unmodelled lower region level with the mCP layer. The same is evident in PBCV-1 (Supplementary Fig. 2e). The ASFV penton protein (Supplementary Fig. 2f) fits the single jelly roll in the same position and orientation as the PBCV-1 penton, with the insertion domain on the outer face of the capsid in density, which neither PBCV-1 nor tokyovirus possess clear density. A BLAST search of the tokyovirus genome, using both the PBCV-1 penton and the Cafeteria-dependent mavirus penton which was modelled into the ASFV penton structure, did not yield any matches, so identification and homology modelling of the penton protein has not been possible. Given the low BLAST metrics for the tokyovirus MCP against both PBCV-1 and mavirus (Table 2) this is not necessarily surprising. To clarify the nature of the protein occupying the lower region of the penton, we must achieve a higher resolution reconstruction.

Immediately around this small pentamer is a symmetric array of further proteins (PC-α, β and γ) within the mCP layer, which act as a replacement for the lattice proteins that support the MCP array in the pentasymmetron and interact with the inner membrane extrusion (Fig. 6d). They may be roughly analogous to the “lantern” proteins in ASFV^19^. These further extend to the edge of the pentasymmetron and interact with the scaffold protein array (Fig. 5). This is a novel structure in NCLDV and plays a role of the formation of the inner membrane extrusion.

In conclusion, tokyovirus shows a departure from the structural features so far demonstrated in published NCLDV cryo-EM studies. While it shares many similarities to the structures of PBCV-1 and ASFV regarding MCP structure, the mCP array is more complex and the free-floating “scaffold” proteins have not been reported previously. The penton protein of tokyovirus is comparable to that of PBCV-1, the closest sequence homolog, with the crystal structure of the penton of another NCLDV fitting neatly. The NCLDVs show a great degree of structural variability in capsid structure while simultaneously demonstrating remarkable similarity in MCP structure.

## Methods

### Tokyovirus growth, purification, and sample preparation

Tokyovirus was propagated in *Acanthamoeba castellanii* cells cultured in PYG medium (2% w/v proteose peptone, 0.1% w/v yeast extract, 4 mM MgSO_4_, 0.4 mM CaCl_2_, 0.05 mM Fe(NH_4_)_2_(SO_4_)_2_, 2.5 mM Na_2_HPO_4_, 2.5 mM KH_2_PO_4_, 100 mM sucrose, pH 6.5). Viruses were purified as described previously^8, 53^. Summarily, the infected culture fluid was collected and centrifuged for 10 min at 1,500 g, 4°C to remove dead cells, before the supernatant was centrifuged for 35 min at 10,000 g, 4°C. The pellet was suspended in 1ml of PBS buffer and loaded onto a 10-60% sucrose gradient, before further centrifugation for 90 min at 8,000 g, 4°C. The concentrated band was extracted and dialysed in PBS before a further round of centrifugation with the same conditions. The pellet was suspended in PBS before cryo-EM grids were prepared.

### Performance Test of 1 MV cryo-HVEM

A JEOL JEM-1000EES cryo-HVEM (JEOL Inc., Tokyo) has been installed into the research center for ultra-HVEM at Osaka University. The microscope is equipped with a LaB^6^ filament electron gun at an accelerating voltage of 1 MV, an autoloader stage can keep up to 12 frozen-hydrated EM grids, and K2 IS direct detector camera (Gatan Inc. USA). The stage and storage of the sample is always cooled with liquid nitrogen that is automatically replenished. Image data are collected manually. For resolution limit test of the 1 MV cryo-HVEM, Pt-Ir film (JEOL Inc., Tokyo) was imaged at a nominal magnification of 100,000× (0.22 Å/pixel on specimen) and 0.6-2 µm defocus. Movie images were recorded on K2 IS camera in counting mode at a dose rate of ∼8 e^-^/pixel/s with 0.2 s/frame for 1 s exposure. Motion-corrected full frames were summed with DigitalMicrograph software (Gatan Inc). For image quality test, MTF and DQE curves were measured using the shadow of a beam stopper metal blade in both counting mode and super-resolution mode of the Gatan K2 IS in cryo-HVEM. The data was processed with FindDQE^54^. For comparison, the data was collected using 300 kV Titan Krios G3 (Thermo Fisher Scientific) and K2 Summit camera with the same conditions.

### Cryo-HVEM data acquisition

An aliquot (2.5 µL) of the purified tokyovirus particles was placed onto R 1.2/1.3 Quantifoil grids (Quantifoil Micro Tools) that were glow-discharged using a plasma ion bombarder (PIB-10, Vacuum Device Inc.) immediately beforehand. This grid was then blotted (blot time: 10 s, blot force: 10) and plunge-frozen using a Vitrobot Mark IV (Thermo Fisher Scientific) with the setting of 95% humidity and 4°C. A total of 304 micrographs were manually collected using a JEOL JEM-1000EES (JEOL Inc., Tokyo) equipped with an autoloader stage and K2 Summit camera (Gatan Inc. USA) optimised for 1 MV HVEM in a total of seven sessions using three grids. Micrograph movie frames were collected in super resolution mode at a magnification equivalent to 1.456 Å/pixel with a target defocus of 2-4 µm. Each exposure was 32 seconds, at a frame interval of 0.2 seconds for a total of 160 frames per micrograph. The frames were stacked using EMAN2^55^ and imported into a beta build of RELION 3.1, before motion correction with the RELION implementation^48^ of MotionCor2^47^. Motion correction was performed using 5×5 patches with B-factor blurring of 500 Å^2^.

### Image processing

Contrast Transfer Function (CTF) estimation of the images was carried out using CTFFIND^56^ 4.1.10 with the following parameters: lower defocus limit, 2500Å; upper defocus limit, 80,000Å; step size, 100; exhaustive search. After the first run using a box size of 512, good CTF fits were selected, and failed fits were re-run using a larger FFT box size. This was repeated until an FFT box size of 2,048. After this point, micrographs which failed to find a good fit were discarded. This resulted in a total of 156 micrographs being carried forward. 1,529 particles were manually picked, extracted with 4× downsampling to 5.824 Å/pixel (2400×2400 pixel boxes become 600×600 pixel boxes) and 2D classified into 40 classes with a circular mask diameter of 2600Å, angular sampling of 2° (automatically increased to 3.75° due to use of GPU acceleration) and a search range of 7 pixels. 1,458 particles were carried over in good classes, and 2D classified a second time with 1° angular sampling (automatically increased to 1.825°) and “ignore CTF until first peak” enabled, resulting in 1,419 particles in good classes. These particles were passed to 3D classification into 5 classes, using 1.8° angular sampling for 25 iterations, followed by 0.5° angular sampling for a further 25 iterations. The two best classes were selected, totalling 1,297 particles. 3D refinement was carried out using only a spherical mask. Particles were re-extracted and re-centred with 3× downsampling (4.368 Å/pixel) and refined further, then re-extracted and re-centred with 2× downsampling (2.912 Å/pixel). Magnification anisotropy refinement and CTF refinement were carried out, improving attained resolution. 3D classification into 5 classes with alignment disabled was carried out using a mask blocking the disordered internal volume (viral DNA) and the best class selected comprising 1,182 particles. Particle polishing had a negligible effect when used. Final post-process resolution depends heavily on the mask used: a capsid-only mask results in a 7.7 Å estimated resolution, including the scaffold proteins and inner membrane lowers this to 8.7 Å. A soft spherical mask gives a 9.4 Å resolution. Local resolution was calculated using the *blocres* module of Bsoft^49, 50^ with no mask applied. The full pathway through the SPA 3D reconstruction is summarized in Supplementary Fig. 6.

### Bioinformatics, model building and fitting

The MCP protein of tokyovirus was identified from the previously reported genome^9^. The primary amino acid sequence of tokyovirus MCP was compared to those of other NCLDVs by BLASTP^57^ and the SWISS-MODEL^40^ server was used to generate a homology model using PDBID: 5TIP^58^ as a template. This homology model was rigid body fit into a tokyovirus MCP extracted from the threefold symmetry axis and sections of the model which lay outside of density were adjusted by COOT^41^ and PHENIX^42^. The central point of the trisymmetron was identified visually and extracted using the UCSF Chimera^59, 60^ Volume Eraser tool, then the volume was cropped to 100^3^ pixels. The resulting model was saved. The trisymmetron map was segmented using the SEGGER^61^ module of USCF Chimera and segments which overlapped the model saved into a separate volume. For the penton protein, the PDB structure of the penton of Cafeteria-dependent mavirus^62^ was fitted to the centre of the five-fold axis and SEGGER was used to extract the region. This also selected the majority of the five contacting MCPs so *molmap* was used to generate a 15 Å model which was passed to the *relion_mask_create* module of RELION 3.1 to generate a binary mask with an extension of 5 pixels and a soft edge of 5 pixels, which was then imposed on the extracted region.

### Visualisation

Data were visualised using the RELION 3.1^48, 63, 64^ *relion_display* module or UCSF Chimera^60^ depending on dimensionality. Pt-Ir power spectra were visualised using Fiji^65^ and rotational averages calculated using the Radial Profile Extended plugin^66^.

## Data availability

Cryo-HVEM reconstructions have been uploaded to the EMDB and are available at the following accession codes, 30797: capsid-only post-process mask, 7.7 Å, 30798: capsid, scaffold and inner membrane post-process mask, 8.7 Å, 30799; soft spherical post-process mask, 9.4 Å.

## Acknowledgements

We thank Masaharu Takemura for his critical reading and valuable suggestions, and Yoko Kayama for initial collection of cryo-EM data. This work was supported by MEXT KAKENHI (JP19H04845 to K.Mu.), the Joint Research of ExCELLS (20-004 to K.Mu.), the Cooperative Study Program of National Institute for Physiological Sciences (20-239 to M.T.), SOKENDAI (to A.C.), Vetenskapsrådet (VR)/The Swedish Research Council (2018-03387 to K.O.), the Swedish Foundation for International Cooperation in Research and Higher Education (STINT) (JA2014-5721 to Janos Hajdu and K.O.), FORMAS research grant from the Swedish Research Council for Environment, Agricultural Sciences, and Spatial Planning (2018-00421 to K.O.), the Royal Swedish Academy of Sciences (BS2018-0053 to K.O.).

## Author contributions

K.Mu conceived the project; K.O. provided the tokyovirus sample. C.S. and K.Mu. prepared cryoEM samples. A.C., R.N.B.S., C.S., N.K., K.Mi., and K.Mu. collected cryo-EM data. N.K. and K.Mi. provided the HVEM performance data. A.C., R.N.B.S., C.S., and K.Mu. processed the cryo-EM data. A.C., R.N.B.S., C.S., and K.Mu. prepared figures and wrote the main manuscript text. All authors reviewed the manuscript.

