## Supplementary INformation for "A novel capsid protein network allows the characteristic inner membrane structure of *Marseilleviridae* giant viruses"

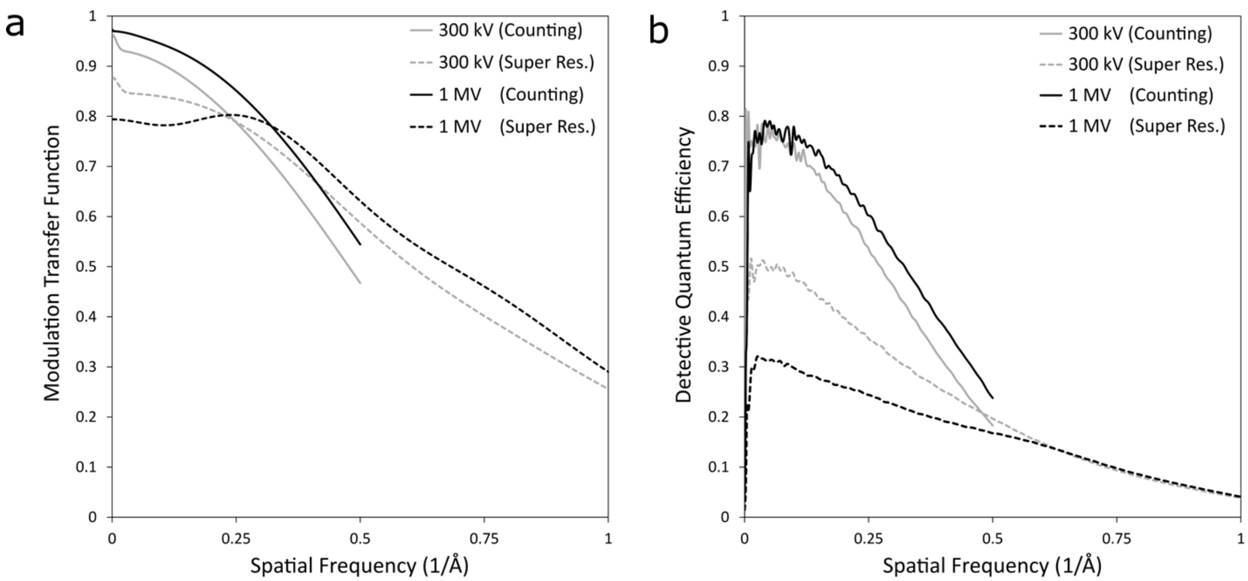


**Fig. S1** Performance of the detector depending on accelerating voltage. a) Modulation transfer function (MTF) curves for the cryo-HVEM (JEOL JEM-1000EES) equipped with a K2 Summit direct electron detector (Gatan Inc.) using electron counting and super resolution modes. 300 kV curves included for comparison, using a Titan Krios G3 (Thermo Fisher Scientific) equipped with a K2 Summit direct electron detector (Gatan Inc.). b) Detective quantum efficiency (DQE) curves for the same conditions as (a).

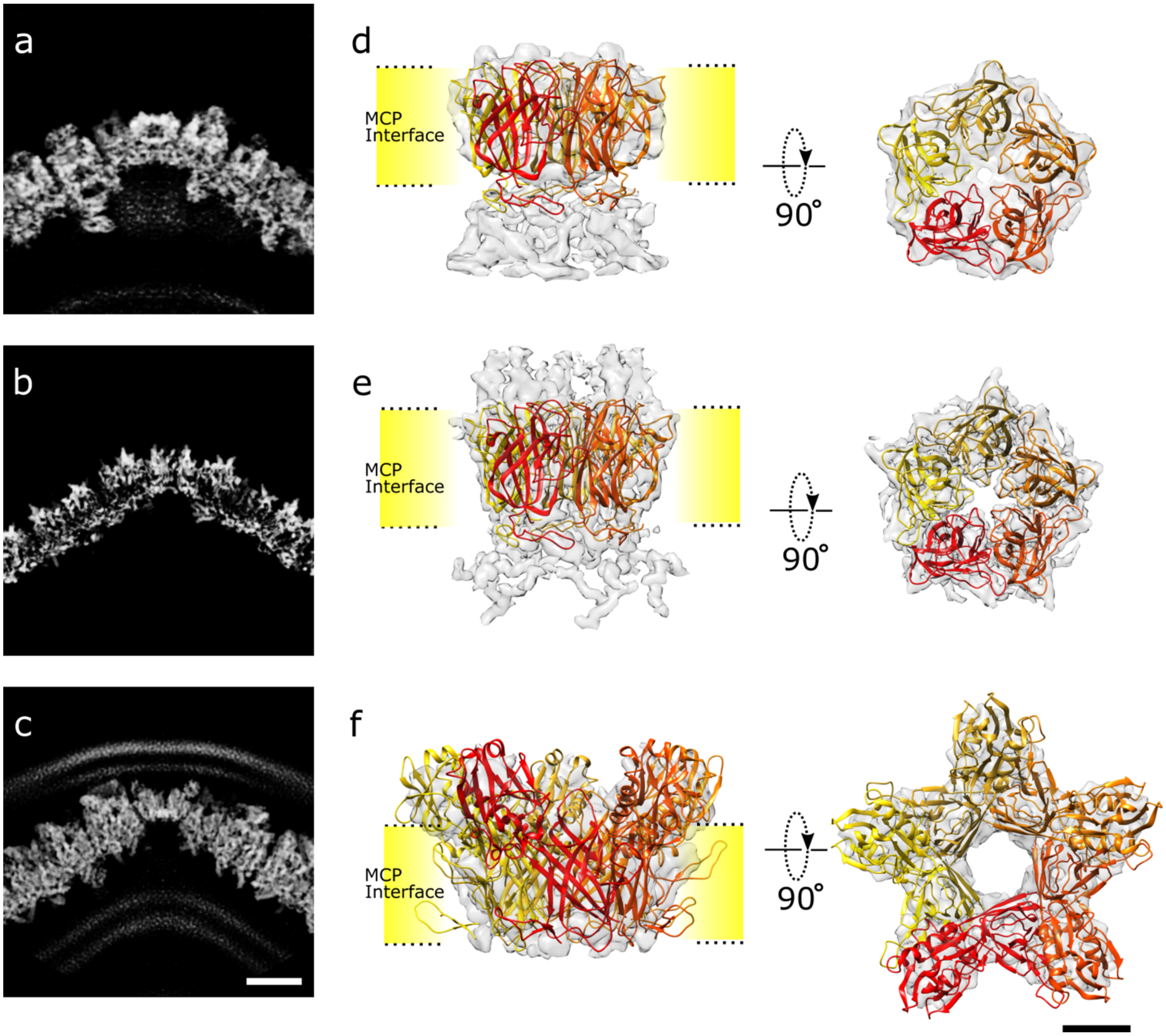


**Fig. S2** Evaluation of penton protein structure. Comparing the slab view of tokyovirus pentasymmetron including penton protein (a) with that of PBCV-1 (b) and ASFV (c). (d) Fitting of the PBCV-1 penton structure (PDBID: 6NCL) to the extracted penton volume of tokyovirus. (e) PBCV-1 penton structure (PDBID: 6NCL) fitted to the PBCV-1 penton cryo-EM map (EMD-0436). (f) ASVF cryo-EM map (EMD-0815) with the Cafeteria-dependent mavirus penton crystal structure (PDBID: 6G41) fitted and modified. Scale bars equal 10 nm in slab views and 2 nm in isosurface/model views.

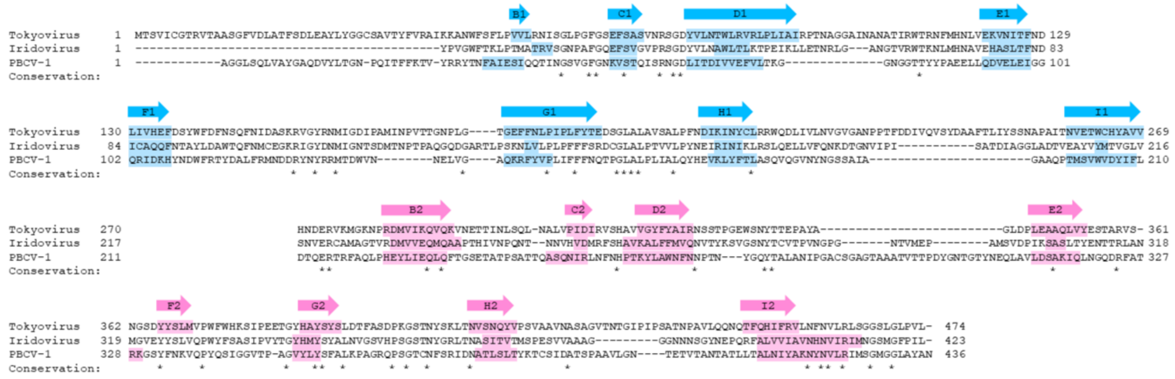


**Fig. S3** Sequence alignment of Tokyovirus MCP versus iridovirus and PBCV-1. Putative β-sheets (coloured arrows) were highlighted. Residues conserved between all three sequences are marked with asterisks.

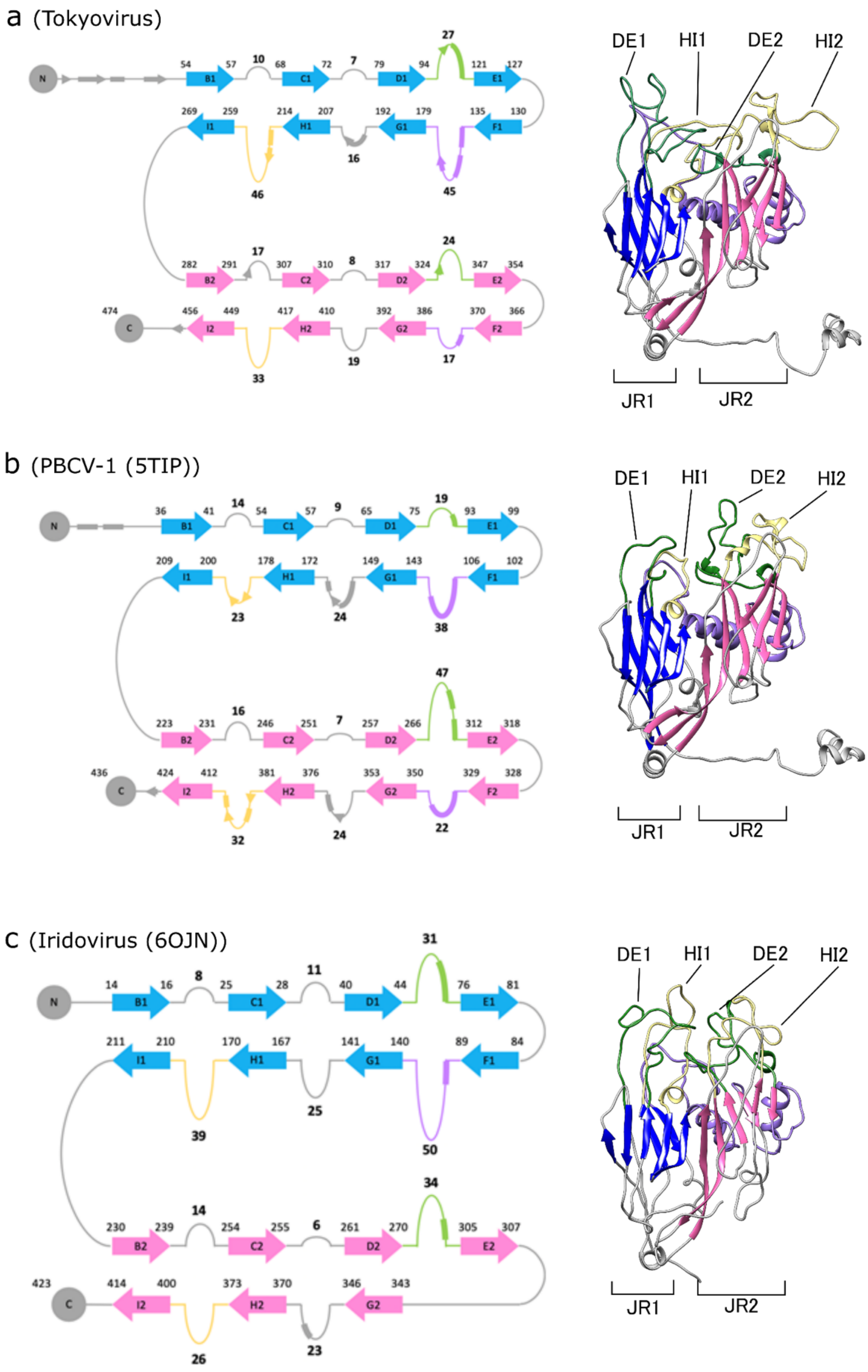
a (Tokyovirus)
N
B1
C1
D1
E1
F1
G1
H1
I1
B2
C2
D2
E2
F2
G2
H2
I2
C
DE1
HI1
DE2
HI2
JR1
JR2
b (PBCV-1 (5TIP))
N
B1
C1
D1
E1
F1
G1
H1
I1
B2
C2
D2
E2
F2
G2
H2
I2
C
DE1
HI1
DE2
HI2
JR1
JR2
c (Iridovirus (6OJN))
N
B1
C1
D1
E1
F1
G1
H1
I1
B2
C2
D2
E2
G2
H2
I2
C
DE1
HI1
DE2
HI2
JR1
JR2

22

**Fig. S4** Comparing the secondary structure of MCP monomers. a) Tokyovirus, b) PBCV-1, c) Iridovirus. For each, jelly roll motif 1 is coloured in blue, jelly roll motif 2 is coloured in pink, DE1 and DE2 loops are coloured in green, FG1 and FG2 loops are coloured in mauve, HI1 and HI2 loops are coloured in pale yellow, other regions coloured in grey.

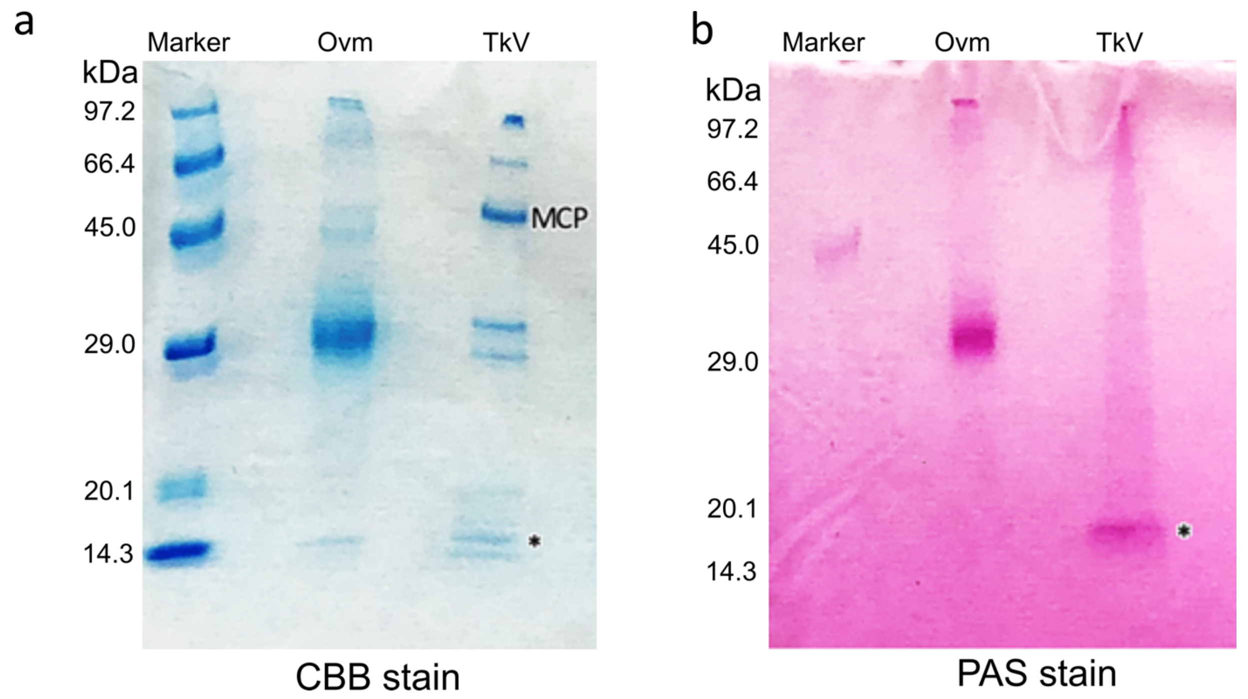


**Fig. S5** Identification of potential glycoproteins by Periodic acid Schiff (PAS) staining. a) SDS-PAGE gel of a positive control, albumin (Ovm), purified tokyovirus (TkV), with strong bands for TkV MCP indicated at ~52 kDa. The gel was stained with Coomassie Brilliant Blue (CBB). b) PAS stained gel with identical loading to that of (a). In the PAS stained gel, no signal is detected at the same MW as the MCP. A single band is identified by PAS staining for TkV at ~14 kDa (marked with asterisks on both gels). The molecular weights of each marker component is indicated on the left of each gel.

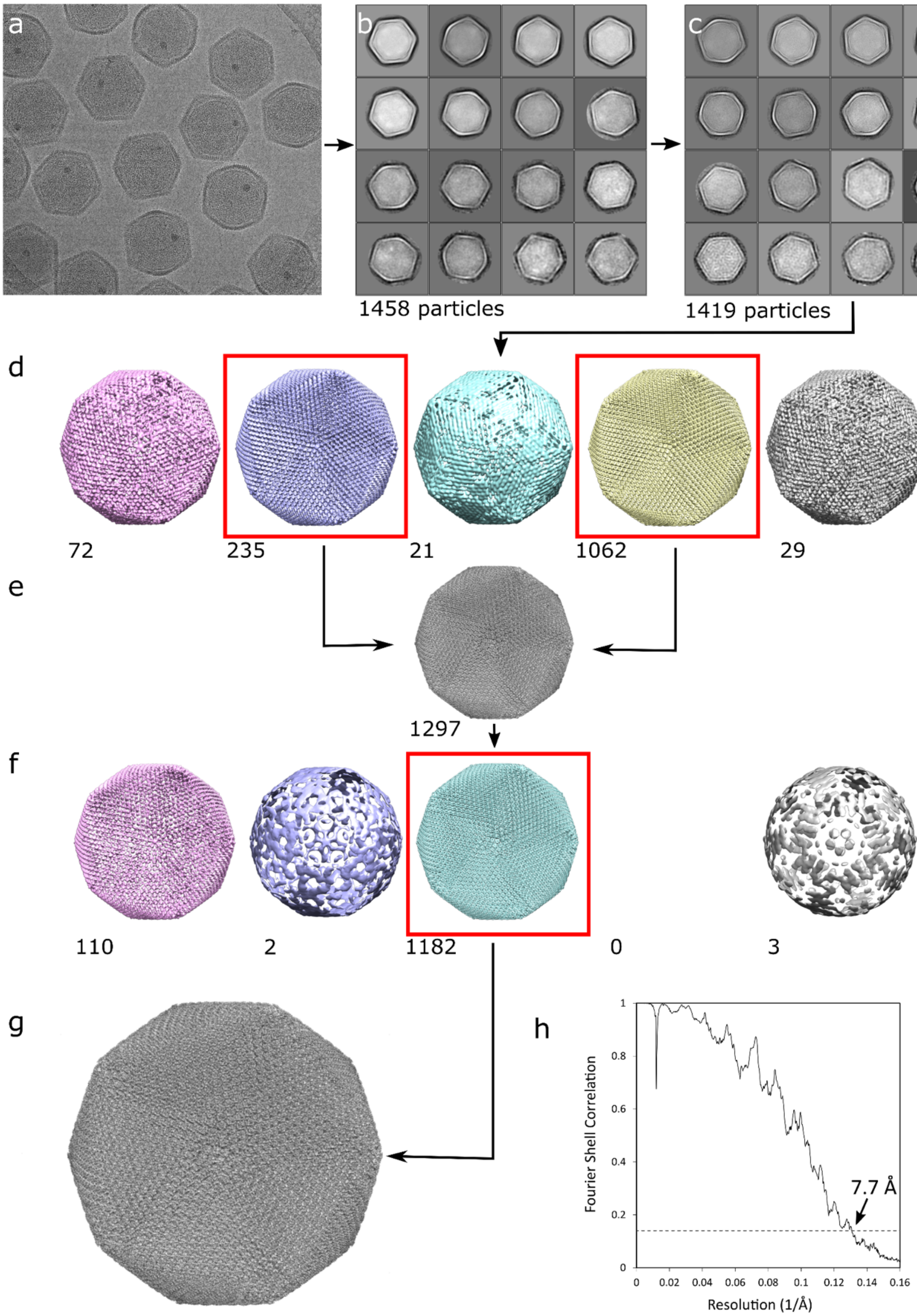
a
b
c
1458 particles
1419 particles
d
72
235
21
1062
29
e
1297
f
110
2
1182
0
3
g
h
Fourier Shell Correlation
Resolution (1/Å)
7.7 Å
0
0.2
0.4
0.6
0.8
1
0.02
0.04
0.06
0.08
0.1
0.12
0.14
0.16

37

**Fig. S6** The pathway through SPA 3D reconstruction of tokyovirus using RELION 3.1. a) A representative micrograph. b) First round of 2D classification. c) Second round of 2D classification. d) 3D classification into five classes with the clearest two chosen for initial 3D refinement. e) Initial 3D refinement. f) After CTF refinement cycles, the reconstruction was again classified into five classes, with the highest resolution one chosen for final 3D refinement. g) Final reconstruction. Numbers by a 3D class indicate particle count. A red box indicates class brought forward through processing. h) Gold standard FSC of the final reconstruction.
